# Movement sonification during haptic exploration shifts emotional outcome without altering texture perception

**DOI:** 10.1101/2025.05.28.656569

**Authors:** Laurence Mouchnino, Pierre-Henri Cornuault, Jenny Faucheu, Arnaud Witt, Chloé Sutter, Benjamin Weiland, Jean Blouin, Francesco Massi, Eric Chatelet, Jérémy Danna

## Abstract

Adding movement sonification to haptic exploration can change the perceptual outcome of a textured surface through multisensory processing. We hypothesized that auditory-evoked emotions would influence the appraisal of textured surfaces, with corresponding changes reflected in cortical excitability. Participants actively rubbed two different textured surfaces (slippery and rough) either without movement sonification, or with pleasant or disagreeable movement sonification. We found that sounds, whether agreeable or disagreeable, did not change the texture appraisal. However, the less pleasant surface was associated with a stronger negative hedonic valence, particularly when paired with disagreeable movement sonification. Time frequency analyses of EEG activities revealed a significant reduction in beta-band power [15-25 Hz] within the source-estimated sensorimotor and superior posterior parietal cortices when contrasting both pleasant and unpleasant sound with the silent touch. This suggests that the primary somatosensory cortices together with the superior parietal regions participated in the audio-tactile binding, with both pleasant and unpleasant sounds. In addition, we observed a significant increase in beta-band power in medial visual areas, specifically when disagreeable movement sonification was paired with tactile exploration. This may reflect a disengagement of visual cortical processing, potentially amplifying auditory-driven emotional responses and intensifying the perceived unpleasantness of the explored surfaces.

## Introduction

Haptic exploration for extracting information about objects is generally considered a silent process. When vision is absent, the texture of an object is thus perceived through multisensory processing of tactile and proprioceptive information (i.e., haptic information). Interestingly, when haptic exploration becomes audible, the perception of texture can change. One of the most compelling demonstrations of tactile-auditory integration altering texture perception is the so-called parchment-skin illusion (Champoux et al. 2011; Jousmäki & Hari, 1998). Specifically, people experiencing this illusion perceive the palms of their hands as becoming either dry or moist, depending on the spectral content of the hand-friction sound played while rubbing hands together. Thus, adding movement-related sounds, a technique called movement sonification, can significantly alter the perceived texture, even though the tactile and proprioceptive inputs evoked during surface exploration remain unchanged. The primary objective of this technique, which is conceptually like the Foley effect in cinema, is to convey movement-related information and enhance both the perception and control of movement (Effenberg, 2005). Applied to pen movements, this technique has been successfully used to improve handwriting performance (Veron-Delor et al., 2020; for a review, see Danna & Velay, 2017). In texture discrimination tasks, haptic exploration has been shown to change in the presence of textured sounds compared to non-textured sounds, notably by increasing movement velocity (Landelle et al., 2021).

A fundamental property of sound is its capacity to evoke a wide range of emotions (Mithen 2009; Dubé & Le Bel 2003). Despite this, little is known about how auditory-evoked emotions influence the appraisal of textured surfaces scanned by a finger. This gap persists although there is well-established evidence that our appreciation of objects is strongly influenced by the tactile sensations and emotions elicited when touching them. The present study aims to determine whether movement sonification during tactile exploration of a surface alters both the perceptual appraisal of the texture and, its emotional valence (ranging from “positive, pleasure” to “negative, displeasure”).

Changes in the perceptual and emotional appraisal of a texture can result from bottom-up and top-down mechanisms. From a bottom-up perspective, the Bayesian approach proposes that the central nervous system integrates information from different sensory modalities based on the estimated probability that it originates from a common or independent source (Körding et al., 2007). The mechanisms underlying cross-modal interactions in the human brain remain largely unknown. This is particularly the case for audio-tactile interactions, despite their frequent occurrence in everyday life, for example, when hearing footsteps on a gravel path. This example complies with the ‘‘temporal principle’’ of multisensory integration stating that an interaction is most likely to be maximally effective when stimuli overlap in time (Meredith et al., 1987). Sound and touch are likely to respond to this principle as they are deeply entangled. Mechanical forces generated by physical contact not only activate somatosensory mechanoreceptors but also generate acoustic waves detectable by the auditory system (Békésy 1959). To date, neurons responsive to both audio and tactile stimuli have been found in primates in the ventral premotor cortex, the ventral intraparietal region and the superior temporal sulcus (e.g., Graziano et al., 1999; Schlack et al., 2005). Further evidence indicates that audio-tactile interactions occur already at early stages of sensory processing. Notably, the somatosensory (S1) and auditory (A1) cortical regions, presumptive unimodal sensory areas, exhibit enhanced responses to bimodal stimuli relative to unimodal inputs (e.g., Landelle et al., 2023). This has been evidenced in animal studies by Lakatos et al. (2007) showing that somatosensory inputs reset the phase of ongoing neuronal oscillations, so that accompanying auditory inputs arrive during an ideal, high-excitability phase, and produce amplified neuronal responses. In contrast, responses to auditory inputs arriving during the opposing low-excitability phase tend to be suppressed. Zhang et al. (2020) extended these findings from primary cortical areas to secondary somatosensory cortice (S2). They showed in the mouse neocortex that, pairing sound with whisker stimulation modulates tactile responses in both S1 and S2. Sound responses of many sound-selective neurons in S2 neurons were spatially colocalized with S2 touch-selective neurons.

Complementing these bottom-up mechanisms, other approaches propose that the perception of a texture under our skin is not driven solely by somatosensory stimulation (see Engel et al., 2001, for a review). Rather, somatosensory processing would be under the influence of top-down control, which actively modulates, and routes sensory inputs based on expectation, attention, and the perceptual task (Talsma et al., 2010; Gilbert et al. 2007 for review). For example, pleasure induced by listening to pleasant music has been shown to require higher levels of attention compared to neutral music, thereby reducing the attentional resources available for other tasks (Nemati et al., 2019). This relationship between attention and the experience of pleasurable sounds can influence how sensory information is processed and integrated, as attentional states have been shown to modulate somatosensory processing (Wiesman & Wilson, 2020).

Through top-down and bottom-up mechanisms, movement sonification may then modulate the hedonic valence of textures during haptic exploration, particularly when the associated sounds convey emotional attributes. In the present study, we hypothesized that texture exploration producing pleasant sounds though movement sonification will be rated as more pleasant, while textures associated with unpleasant sounds will be perceived as less pleasant. These perceptual effects of movement sonification may be linked to changes in beta power within cortical areas involved in processing somatosensory inputs (e.g., S1, PPC) as well as in areas engaged in evaluating emotional valence, such as the cingulate cortex.

## Methods

### Participants

Twelve right-handed adults (6 women, 6 men, mean age 26 ± 2), without any known neurological or motor disorders, participated in the experiment. All participants gave their written informed consent to take part in this study, which conformed to the standards set in the Declaration of Helsinki. This study was approved by the local Ethics Committee (CERSTAPS: IRB00012476-2021-09-12-140).

### Task

The participants were asked to stroke their finger across the sample surface in a disto-proximal direction several times (∼40 times over 4 recording sessions of 15 seconds), lifting the finger between each stroke. They used their dominant hand and had their eyes closed. Sliding velocity, contact load and finger/surface angle were freely chosen by the participants. After completing the series of strokes, participants indicated whether they felt the texture they just explored as slippery/vibrating/rough (perceptual outcome) and rated their liking of the texture using a 4-levels hedonic scale (emotional outcome): I like it a lot / I like it / I don’t like it / I really don’t like it.

Two textured panels made of polyurethane resin (P1 and P2, 60 × 26 mm) were selected according to the results by Faucheu et al. (2019). In their study, among the 51 explored textures, P1 was clearly qualified as SLIPPERY and P2 as ROUGH. In addition, the hedonic rating of the panels was consensual across the 43 participants: P1 was rated as likeable, and P2 as disagreeable. For both panels, the texture consisted of a flat surface patterned with cylindrical dots, evenly arranged in a hexagonal array. The panels varied in height, diameter and spacing between the cylindrical dots (see Table 1 and Fig. 1A, left panel).

**Table 1.** Topographic characteristics of P1 and P2 panels.

| P1 AND P2 PANELS | Height ($\mu\text{m}$) | Diameter ($\mu\text{m}$) | Inter-dot spacing ($\mu\text{m}$) |
| --- | --- | --- | --- |
| P1 | 19 | 22 | 28 |
| P2 | 18 | 106 | 610 |

**Figure 1.**
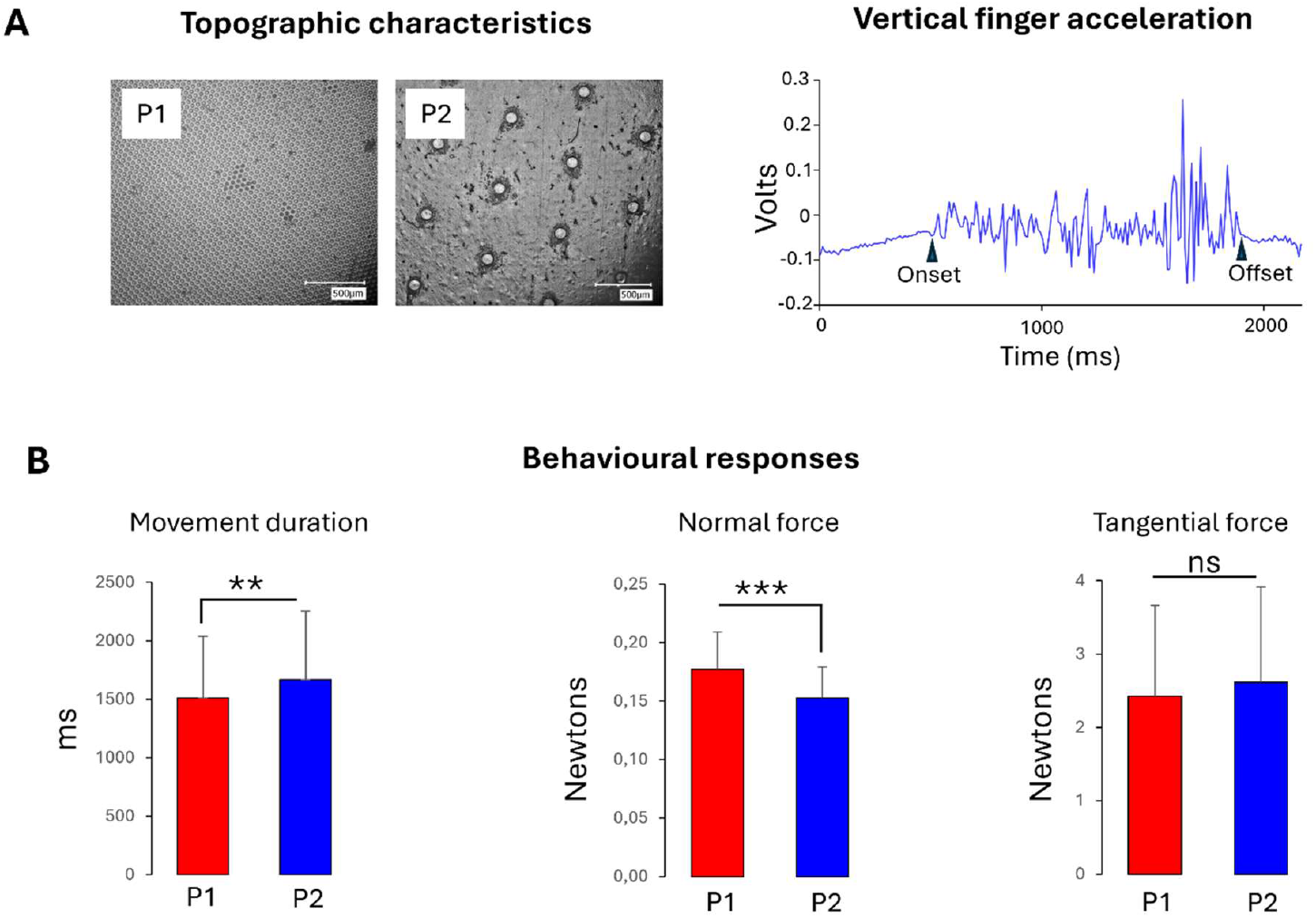
Microscale topographic views of the slippery and rough surfaces (left panel) and example of vertical acceleration recorded during a single sliding finger movement across the P2 rough surface (right panel) (**A**) and behavioral data: movement duration, normal and tangential forces exerted on the surfaces (respectively left, middle and right panels) **(B**).

### Movement sonification

Real time sonification of the finger movement was ensured by an optical motion tracking system fixed to fingernail (sampling frequency: 100 Hz), using a Max/MSP® program. The musical sonification was based on a mapping between the finger’s positions and musical chords with or without the presence of false notes (e.g. audio recordings of the false notes are available at: https://www.gdr.tact.uha.fr/groupes-de-travail/comprendre-et-caracteriser/). The sonified movement was revealed by loudspeakers placed in front of the finger as it moves across the surface. The musical chord was tested on some fifteen individuals prior to the experiment. The sonification without false notes was deemed agreeable, while the sonification with false notes was considered disagreeable by most participants.

### Recordings and analyses

To record normal and tangential (shear) force data from the finger/surface interaction, the textured surface is mounted on the top of a home-made multi-sensors box comprising three resistive normal load sensors and one piezoelectric force sensor (Kistler 9217A). Data are acquired through a National Instrument data acquisition system (NI PCIe-6321) using a 10 kHz sampling frequency, and recorded with Labview software.

The vertical acceleration of the right index finger was recorded at 1000 Hz with a small and light (5 mm x 12 mm x 7 mm, 0.8 g) accelerometer (PCB 352A24, PCB Piezotronics, Inc., Depew, USA) secured on the finger’s nail with wax. With a high sensitivity of 100 mV/g and a measurement range of ± 50 g pk, this accelerometer can detect very low levels of acceleration. The onset and offset of the finger acceleration were used to define as the onset and offset of the stroking movement, respectively (Fig1A, right panel).

Electroencephalography (EEG) activity was continuously recorded from 64 Ag/AgCl surface electrodes embedded in an elastic cap (BioSemi ActiveTwo system: BioSemi, Netherlands). Specific to the BioSemi system, “ground” electrodes were replaced by Common Mode Sense active and Driven Right Leg passive electrodes. The signals were pre-amplified at the electrode sites, post amplified with DC amplifiers, and digitized at a sampling rate of 1024 Hz (Actiview acquisition program). The signals of each electrode were referenced to the mean signal of all electrodes. The continuous EEG signal was segmented into epochs aligned with the onset of the monotonic increase in shear forces generated by the finger on the surface. The average amplitude computed 100 ms prior to the force onset served as baseline. A mean of 35 trials for P1 and 32 trials for P2 were included in the analyses; the number of trials did not significantly differ between the 3 conditions of sonification (F_2,18_ = 3.1; *p* > 0.05).

### Cortical sources

Neural sources were estimated with the dynamical Statistical Parametric Mapping (dSPM, Dale et al., 2000) implemented in the Brainstorm software. A boundary element model (BEM) with three realistic layers (scalp, inner skull, and outer skull) was used to compute the forward model on the anatomical MRI brain template from the Montreal Neurological Institute (MNI Colin27). Using a realistic model has been shown to provide more accurate solution than a simple three concentric spheres model (Sohrabpour et al., 2015). We used of a high number of vertices (i.e., 15 002 vertices) to enhance the spatial resolution of the brain template. Such EEG source reconstruction has proved to be suited for investigating the activity of outer and inner cortical surfaces with 64 sensors (Chand and Dhamala, 2017; Ponz et al., 2014). Measuring and modelling the noise contaminating the data is beneficial to source estimation. Noise covariance matrices were computed using the trials recorded while the index finger stayed stationary on the surface.

Beta band (15-25 Hz) event-related synchronization / desynchronization (ERS/ERD) was computed relative to a baseline window from - 3000 to -500 ms prior to finger movements and then averaged over 1000 ms interval starting at the onset of the strokes. This approach ensured that our ERD/ERS analyses captured most of the exploratory movement, encompassing approximately two-thirds of its total duration.

### Statistics

The normal distribution of data sets was assessed using the Kolmogorov–Smirnov test to validate the use of parametric tests. For the behavioral data, analyses of variance (ANOVAs) were conducted on movement duration, normal force and tangential force using the following design: Surface (2 types of surfaces: P1 and P2) × Sonification (3 modalities: silent, agreeable, and disagreeable). For the categorical data on perceptual and emotional outcomes, Pearson chi-square (χ^2^) tests were conducted to compare the two texture panels (P1 and P2) and the three sonification conditions. The EEG data was submitted to paired *t*-tests implemented in Brainstorm software.

## Results

### Behaviour

Non-parametric (χ^2^) tests revealed that the interactions between Surface and Sonification were not significant for any of the behavioral variables: stroke duration (F _2,22_ = 0.39; p = 0.67); normal forces (F _2,22_ = 1.80; p = 0.18) and tangential forces (F _2,22_ = 1.37; p = 0.27) (Fig.1B).

For simplicity and consistency, subsequent analyses will be conducted separately for the factors Surface and Sonification.

Although no specific instructions were given regarding the finger movement, stroke durations were slightly but significantly shorter on the slippery surface (P1: 1512 ± 527 ms) compared to the rough surface (P2: 1668 ± 586 ms), as confirmed by a significant surface effect (F _1,11_ = 13.18; p <0.01). The shorter stroke duration observed on the slippery surface cannot be attributed to smaller shear (tangential) forces as these forces did not differ significantly between both surfaces (F _1,11_ = 0.55; p = 0.47). On the other hand, the normal force exerted by the finger was unexpectedly higher on the P1 slippery surface (0.18 ± 0.3N) than on P2 rough surface (0.15 ± 0.3N) (F _1,11_ = 21.73; p < 0.01).

The forces (normal or tangential) exerted by the finger while rubbing the surfaces were not significantly affected by the movement sonification (F _2,22_ = 3.18; p = 0.061 and F _2,22_ = 3.13; p = 0.063, for the normal and tangential forces, respectively).

### Perceptual outcome

Participants categorized P1 and P2 surfaces differently (χ^2^ = 168,61; *p* < 0.001). P1 was overwhelmingly perceived as slippery (84% on average), whereas P2 was primarily identified as rough, though with less consensus (63% on average) (Fig.2A). Additionally, P2 was also identified as vibrating in about one-third of the responses (34% on average). These surface categorizations were consistent with those reported by Faucheu et al. (2019) using the same materials. The sonification, either agreeable or disagreeable, did not change the texture appraisal (i.e., perceptual outcome) as suggested by the absence of significant effect of the Pearson chi-square on P1 (χ^2^ =5.61; *p* = 0.23) nor on P2 (χ^2^ = 2.73; *p* = 0.60).

**Figure 2.**
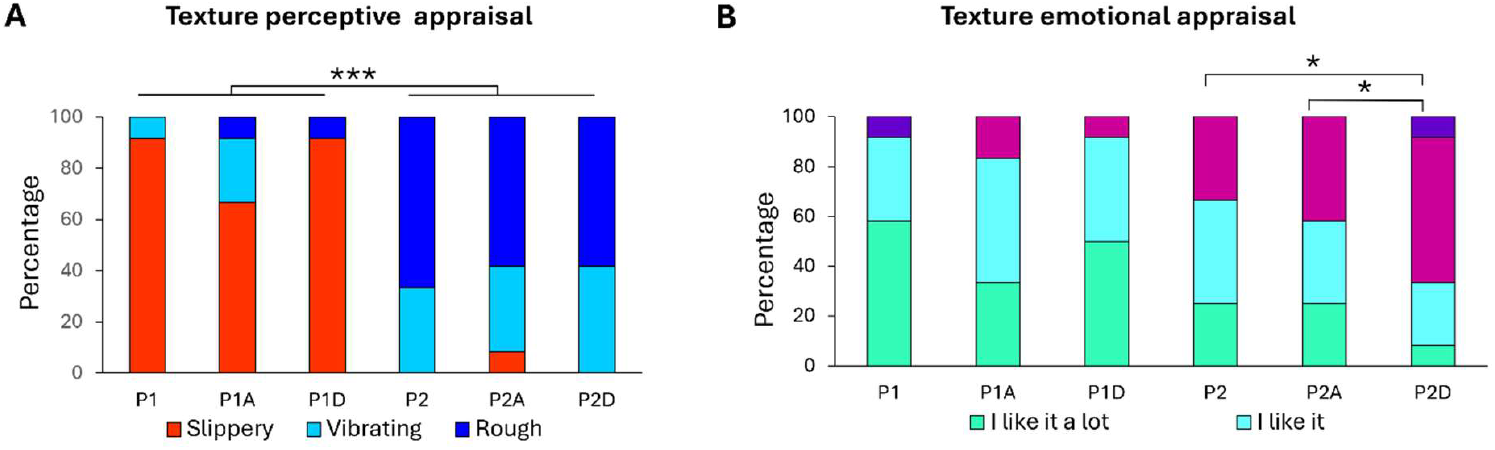
Texture perceptual appraisal (**A**) and. emotional appraisal (**B**) of the P1 slippery and P2 rough surfaces, without stroke sonification, or with either agreeable (P1A, P2A) or disagreeable (P1D, P2D) sonification. (*, p < 0.05 and ***, p < 0.001).

### Emotional outcome

To examine the relationship between movement sonification conditions and surface hedonic judgement, emotional appraisal was evaluated using a Likert scale. For the slippery surface P1, adding either a pleasant or unpleasant sound did not change the hedonic perception of the surface (*p* = 0.67). A striking finding is that only the rough, and less pleasant P2 surface showed a change in its hedonic valence depending on the sonification conditions (χ^2^ = 6.06; *p* = 0.04, Fig.2B). The P2 surface was perceived as more unpleasant with disageeable sonification compared to when it was stroked without movement sonification (*p* = 0.01). The pleasant sonification did not change the emotional output (*p* < 0.05) (Fig.2B).

### Brain activity

Given that the power of beta-band neural oscillations is inversely correlated with cortical processing (Cheyne et al. 2003; Pfurtscheller and Lopes da Dilva, 1999), contrasting the topographical distribution of beta power between both sonification conditions and the silent stroke condition can provide reliable estimate of the effect of movement sonification on cortical processing. For the P2 rough surface, the contrasts revealed a significant reduction in beta-band power within the source-estimated sensorimotor and superior parietal cortices during stroking under both sonification conditions, relative to the silent stroke condition (Fig. 3A, cold blue color). This decreased beta-band power (ERD) suggests amplified neuronal responses when the somatosensory inputs, particularly of tactile origin from the stroking movement, are paired with auditory inputs. However, in other cortical regions, beta-band power varied depending on the valence of the sonification.

**Figure 3.**
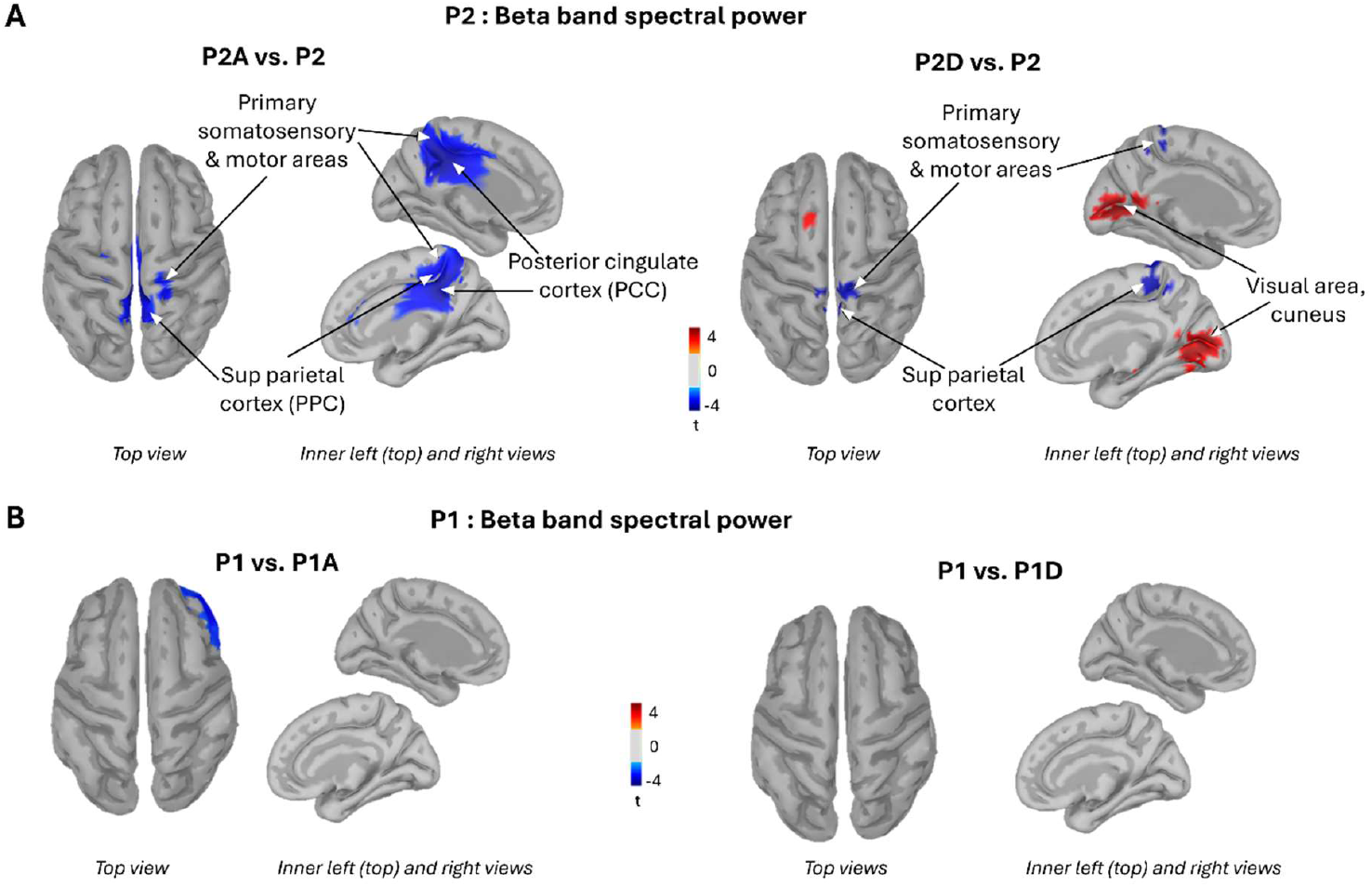
Topographical distribution of beta band power resulting from the contrasts: P2 rough surface with agreeable (left panel) and disagreeable (right panel) sonifications vs P2 with silent stroke **(A)**, P1: slippery surface with agreeable (left panel) and disagreeable (right panel) sonifications vs P1 silent stroke **(B)**.

The agreeable movement sonification elicited an ERD largely circumscribed to the posterior cingulate cortices (Fig. 3A, left panel). In contrast, the disagreeable sonification induced an increased beta-band power (ERS) over both the right and left primary visual cortices relative to the silent condition (indicated by the warm orange color in Fig. 3A, right panel). The valence of the movement sonification had no effect on beta-band power when stroking the P1 slippery surface (Fig. 3B). Overall, pairing an auditory stimulus with finger movement significantly altered cortical processing when participants stroked the rough, unpleasant P2 surface, but had no observable effect during contact with the smooth, pleasant P1 surface.

## Discussion

The aim of this study was to investigate whether movement sonification alters the perceptual evaluation and hedonic valence of textured surfaces, as assessed through behavioural, psychophysical, and neurophysiological measures. The results show that texture appraisal is impervious to sonification conditions, whereas their emotional valence is not. This effect was observed with the P2 rough surface, which was classified as disagreeable and perceived as even more unpleasant when paired with disagreeable movement sonification. However, the brain’s response to the audio-tactile integration was altered by both the agreeable and the disagreeable movement sonifications. In both conditions where stroking movement evoked auditory inputs, we observed a marked decrease in beta-band oscillatory power over the sensorimotor cortex and the superior parietal cortex as compared to silent touch (Fig. 3A, ERD blue/cold color). Such decreases in beta-band power are commonly interpreted as reflecting increased cortical excitability and enhanced sensory processing (Haegens et al. 2011; Romei et al. 2008; Sauseng et al. 2009; Kilavik 2012 for review). It is also in line with the fMRI study by Landelle et al.’s (2023), which demonstrated the sensitivity of the primary somatosensory cortex to tactile input from the hand. Together, these findings suggest that multisensory interactions are not confined to higher-order brain regions but also take place in early somatosensory areas including primary somatosensory cortices and superior parietal regions. This suggestion is supported by neuroanatomical studies showing direct thalamocortical projections to the superior parietal areas in macaques (Pearson et al. 1978; Pons and Kaas 1985; Padberg et al. 2009; Impieri et al. 2018). The early multisensory interaction is thought to be mediated by means of phase resetting. The presentation of a stimulus to one sensory modality (e.g., auditory) resets the phase of ongoing oscillations in another modality (e.g., tactile) such that processing in the latter modality is modulated (Lakatos et al., 2007, 2009).

While these data speak to the importance of primary somatosensory regions for audio-tactile information integration, evaluating the hedonic valence of movement sonification likely engages higher-order brain areas. One candidate region is the posterior cingulate cortices (PCC), which exhibited a significant increase in beta band ERD (15-25 Hz) during agreeable sonification compared to silent touch. However, this change in beta band power was not accompanied by a corresponding change in the emotional rating of the scanned surface. It is possible that altering the hedonic valence of the touched surface would require a sound that is not merely free of dissonance but is explicitly pleasant. The fact that PCC beta-band power remained unchanged during disagreeable sonification suggests that this medial cortical region may be specifically involved in processing pleasantness. In this light, our findings are consistent with studies showing that the PCC is engaged in evaluating the emotional valence (Maddock et al., 2003) and context of sensory information via pathways from somatosensory regions (superior and inferior parietal areas, Vogt, 2014), auditory regions (Vogt & Laureys, 2009) and adjacent cingulate areas such as the midcingulate cortex (Vogt 2016; Rolls, 2014; Grabenhorst & Rolls, 2011). Due to these extensive connections, the PCC is considered to exhibit the highest degree of cross-network interaction (Betti et al., 2021 for review).

When false notes were present, as during the disagreeable movement sonification of P2 rough surface, a bilateral decrease in brain activity was observed in the early visual cortex compared to silent touch. This was evidenced by an increase in event related synchronization (ERS) (red/warm color, Fig. 3A, right panel) in occipital regions (BA17). This observation raises the question as to how and why the earliest visual cortical area, known to receive direct retinal inputs from the thalamic lateral geniculate nucleus, is modulated by auditory signals. Signatures of auditory inputs in primary visual cortex were previously observed by Vetter et al.’s (2014) fMRI study in absence of visual inputs (see also Manson et al. 2019). Their results showed that the early visual cortex contributes, via its corticocortical connections, to the encoding of complex natural sounds as abstract, higher-level internal representations (Petro et al., 2017 for a review), further highlighting the existence of multiple cortical representations of sounds. The disengagement of visual cortical areas during disagreeable movement sonification (reflected by increased beta-band ERS) may have played a crucial role in enhancing auditory-driven emotional responses by reducing the influence of representations unrelated to auditory pleasantness. This mechanism could explain why participants rated the rough surface as even more unpleasant when their haptic exploration was paired with dissonant sonification (see Fig. 2A).

Our results may be interpreted within the framework of the Bayesian causal inference model that postulates that multisensory integration is a consequence of an optimal probabilistic inference of each sensory modality (Kayser & Shams, 2015). Accordingly, both sensory sources, auditory and tactile, are integrated in the judgment task but with a different weight allocated to each sensory input, which depends on the level of relevance and accuracy of the processed information. For instance, for the surface P2, the addition of disagreeable sounds shifts the emotional response towards the unpleasant, as the emotional categorization of the tactile modality is less stabilized as shown by the emotional rating distribution. The lack of effect of sound conditions on P1 slippery agreeable surface on neural excitability (Fig. 3B) suggests that the tactile modality has a greater weight than the auditory modality in this texture perception task at both perceptual and emotional levels. Indeed, adding an unpleasant sound is not decisive when the tactile experience is considered agreeable. This aligns with findings in young adults, who were able to segregate irrelevant auditory information to maintain their haptic discrimination performance (Lederman, 1979; Landelle et al., 2021).

## Acknowledgments

We thank Gilles Marivier and Pauline Couta, society Weydo Music, for creating the sonification set up.

